# Cellular Iron Deficiency Disrupts Thyroid Hormone Regulated Gene Expression in Developing Hippocampal Neurons

**DOI:** 10.1101/2023.06.17.545408

**Authors:** Timothy R. Monko, Emma H. Tripp, Sierra E. Burr, Karina N. Gunderson, Lorene M. Lanier, Michael K. Georgieff, Thomas W. Bastian

**Affiliations:** University of Minnesota, School of Medicine, Department of Pediatrics; University of Minnesota, Department of Neuroscience

**Keywords:** Iron, thyroid hormone, energy metabolism, neuron development, primary neuronal culture

## Abstract

**Background:** Developing neurons have high thyroid hormone and iron requirements to support their metabolism and growth. Early-life iron and thyroid hormone deficiencies are prevalent, often coexist, and increase the risk of permanently impaired neurobehavioral function in children. Early-life dietary iron deficiency reduces thyroid hormone levels and impairs thyroid hormone-responsive gene expression in the neonatal rat brain.

**Objective:** This study determined whether neuronal-specific iron deficiency alters thyroid hormone-regulated gene expression in developing neurons.

**Methods:** Iron deficiency was induced in primary mouse embryonic hippocampal neuron cultures with the iron chelator deferoxamine (DFO) beginning at 3 days in vitro (DIV). At 11DIV and 18DIV, mRNA levels for thyroid hormone-regulated genes indexing thyroid hormone homeostasis (*Hr*, *Crym*, *Dio2*, *Slco1c1*, *Slc16a2*) and neurodevelopment (*Nrgn*, *Pvalb*, *Klf9*) were quantified. To assess the effect of iron repletion, DFO was removed at 14DIV from a subset of DFO-treated cultures and gene expression and ATP levels were quantified at 21DIV.

**Results:** At 11DIV and 18DIV, neuronal iron deficiency decreased *Nrgn, Pvalb,* and *Crym*, and by 18DIV, *Slc16a2, Slco1c1, Dio2,* and *Hr* were increased; collectively suggesting cellular sensing of a functionally abnormal thyroid hormone state. Dimensionality reduction with Principal Component Analysis (PCA) reveals that thyroid hormone homeostatic genes strongly correlate with and predict iron status (*Tfr1* mRNA). Iron repletion from 14–21DIV restored neurodevelopmental genes, but not all thyroid hormone homeostatic genes, and ATP concentrations remained significantly altered. PCA clustering suggests that cultures replete with iron maintain a gene expression signature indicative of previous iron deficiency.

**Conclusions:** These novel findings suggest there is an intracellular mechanism coordinating cellular iron/thyroid hormone activities. We speculate this is a part of homeostatic response to match neuronal energy production and growth signaling for these important metabolic regulators. However, iron deficiency may cause permanent deficits in thyroid hormone-dependent neurodevelopmental processes even after recovery from iron deficiency.

## Introduction

Human brain development is a dynamic, metabolically demanding process, which must occur in a sequential and coordinated fashion. The rapidly growing neonatal human brain accounts for ∼60% of the total body energy utilization, (compared to 20% in the adult brain), indicating the high energy and nutrient requirements of proper brain growth (1, 2). Iron and thyroid hormone are two critical regulators of both systemic and cellular energy production and homeostasis. Thyroid hormone regulates cellular energy (ATP) demand and utilization in the developing brain, through its regulation of genes involved in cellular energy metabolism and in stimulation of ATP-consuming processes, including generation of ion gradients, cytoskeletal dynamics, and DNA replication for cell proliferation (3). Iron regulates cellular energetic capacity in the developing brain through its direct role in the structure and redox activity of cytochrome- and iron-sulfur cluster-containing TCA cycle and electron transport chain proteins (4). Neurons, which rely predominantly on iron- and thyroid hormone-dependent mitochondrial respiration for their energy needs, are most sensitive to perturbations in iron and thyroid hormone during rapid neonatal development. Isolated fetal-neonatal iron deficiency or thyroid hormone deficiency cause similar neurodevelopmental impairments, including less complex neuronal structure, lower electrophysiologic capacity and deficits in learning, memory, psychosocial, and motor skills (5, 6). Deficits persist into adolescence and adulthood despite iron or thyroid hormone repletion, causing life-long brain dysfunction and increased risk for other brain disorders, including autism and schizophrenia (5, 6).

More than 2 billion people worldwide are at risk of developing iron deficiency (with or without anemia) or thyroid hormone deficiency (due to iodine deficiency or genetic disorders), despite widespread efforts to reduce their prevalence (7–10). Thyroid hormones—thyroxine (T4) and triiodothyronine (T3)—are only produced in the thyroid gland via thyroid peroxidase, which requires iron for its enzymatic activity (11). Iron deficiency causes impaired thyroid function in pregnant women and children and often occurs simultaneously with iodine deficiency (12, 13). Moreover, iron deficiency blunts and iron treatment improves the efficacy of iodine prophylaxis on thyroid function in iodine-deficient children.

We, and others, previously demonstrated that fetal-neonatal iron deficiency reduces circulating T4 and T3 concentrations, lowers brain T3 levels, and impairs brain thyroid hormone-responsive gene expression in neonatal rats (14–17), suggesting that iron deficiency-induced systemic- and brain- thyroid hormone deficiency may contribute additional deleterious effects on brain development beyond iron deficiency itself. *De novo* thyroid hormone synthesis only occurs in the thyroid gland and is mediated by the iron-dependent enzyme thyroid peroxidase (11). Thus, the effect of iron deficiency on brain thyroid hormone levels and activity was proposed to be downstream of a direct effect on the thyroid gland and an unfortunate maladaptive consequence for the developing iron-deficient brain. However, it may be advantageous for the developing brain to metabolically match neuronal metabolic and growth rates—mediated by thyroid hormone—with availability of metabolic substrates for oxidative phosphorylation (e.g., iron) to prevent damaging effects of metabolic dyshomeostasis. Whether such neuron-intrinsic coordination of iron and thyroid hormone occurs within the developing brain is unknown.

Intracellular T4/T3 levels and activity are controlled in a tissue- and cell type-specific manner by thyroid hormone transporters (e.g., Oatp1c1/*Slco1c1* and Mct8/*Slc16a2*), deiodinase activity (e.g., D2/*Dio2* and D3/*Dio3*), and intracellular thyroid hormone-binding proteins (e.g., Crym) (3). Thyroid hormone predominantly controls critical brain developmental processes through regulation of gene transcription (3, 18). T3 binds to nuclear thyroid hormone receptors at specific thyroid hormone response elements (TREs) in the upstream promoter region of thyroid hormone-responsive genes. For positively regulated genes, thyroid hormone receptor binding of T3 alters the protein’s conformation and signals for the recruitment of co-activators, resulting in activation of thyroid hormone-responsive gene transcription. In the absence of T3, thyroid hormone receptors bound to the thyroid hormone response element of positively regulated thyroid hormone-responsive genes recruit co-repressors (e.g., Hairless, Hr), resulting in down-regulation of gene transcription. Genes that are negatively regulated by thyroid hormone are typically involved in maintaining thyroid hormone homeostasis (e.g., *Dio2*, *Slc16a2*, *Slco1c1*). Thyroid hormone regulates the expression of numerous genes involved in key neurodevelopmental processes including transcriptional regulation (e.g., *Hr* and *Klf9*), neurotrophic signaling (e.g., *Bdnf*), axon myelination (e.g., *Mbp*), synaptic plasticity (e.g., *Nrgn* and *Pvalb*), and energy metabolism (e.g., *Cox IV*). Iron also regulates many of the same genes (19), providing further support for a co-regulation of iron and thyroid hormone activities in the developing brain.

The objective of the current study was to test the hypothesis that iron control of thyroid hormone activity occurs in a cell-intrinsic manner specifically within the developing neuron. Understanding the relationship between iron deficiency and thyroid hormone gene response may reveal an apparent adaptive thyroid hormone response to changes in iron status. Our mouse embryonic hippocampal neuron culture model of chronic iron deficiency uniquely allows us to test this hypothesis throughout the physiological development of the functionally relevant cell type. This approach dissociates the iron and thyroid hormone requirements of the developing neuron from the iron-dependent *de novo* thyroid hormone synthesis that occurs only in the thyroid gland. Our findings show that neuronal-specific iron deficiency reduces thyroid hormone-regulated gene expression, indicative of a functionally abnormal thyroid hormone status, despite normal thyroid hormone availability. This suggests an undiscovered intracellular mechanism coordinating cellular iron and thyroid hormone activities during neuronal development, potentially as an adaptive response to preserve neuron function in adverse conditions.

## Materials and Methods

### Animals

Timed-pregnant CD1 mice were obtained from Charles River Laboratories (Wilmington, MA). Mice were given free access to food and drinking water and were housed at constant temperature and humidity on a 12h light:dark cycle. All animal procedures were conducted in facilities accredited by the Association for Assessment and Accreditation of Laboratory Animal Care (AAALAC) and in accordance with the principles and procedures outlined in the NIH Guide for the Care and Use of Laboratory Animals. The local Institutional Animal Care and Use Committee approved these procedures.

### Primary hippocampal neuron culture

The experimental design and treatment group designations are outlined in Figure 1. Primary hippocampal cells were collected and pooled from multiple embryonic day 16 mixed-sex mouse embryos and plated in neuronal plating medium (10 mM HEPES, 10 mM sodium pyruvate, 0.5 mM glutamine, 12.5 μM glutamate, 10% fetal bovine serum (FBS), 0.6% glucose in minimal essential medium plus Earl’s salt) at 160cells/mm^2^ in 35 mm dishes for gene expression, 6-well plates for culture media thyroid hormone concentrations, and 96-well plates for culture media thyroid hormone and neuronal ATP analyses as previously described (20–22). After 2 hours, plating medium was removed and switched to neuronal growth medium (Neurobasal containing 2% B27 serum-free supplement, 0.5 mM glutamine and 100 units/mL Penicillin-Streptomycin; Thermo-Fisher), which contains a proprietary concentration of T3 that was not experimentally manipulated (23). Because *de novo* synthesis of thyroid hormones only occurs in the thyroid gland, this means all cultures were performed under thyroid hormone- sufficient conditions. Neuronal growth medium was “conditioned” on separate postnatal mixed- glia cultures. At 3 days *in vitro* (DIV), cultures were treated with 67.5μM 5-fluoro-2’-deoxyuridine (Sigma-Aldrich #F0503; Burlington, MA)/136μM uridine (Sigma-Aldrich #U6381; Burlington, MA) (5-FU), an anti-mitotic drug used to inhibit glia proliferation. Functional neuronal FeD was accomplished by treating cultures with 9μM deferoxamine (DFO, Cayman Chemicals #14595; Ann Arbor, MI) beginning at 3DIV. Iron-sufficient cultures (0µM DFO) were treated in the same manner except with vehicle (sterile, ultrapure water in neuronal growth medium). Each week, beginning at 7DIV, half of the medium was removed and replaced with fresh glia-conditioned neuronal growth medium containing 5-FU and 0µM or 9µM DFO for iron-sufficient or -deficient cultures, respectively.

**Figure 1.**
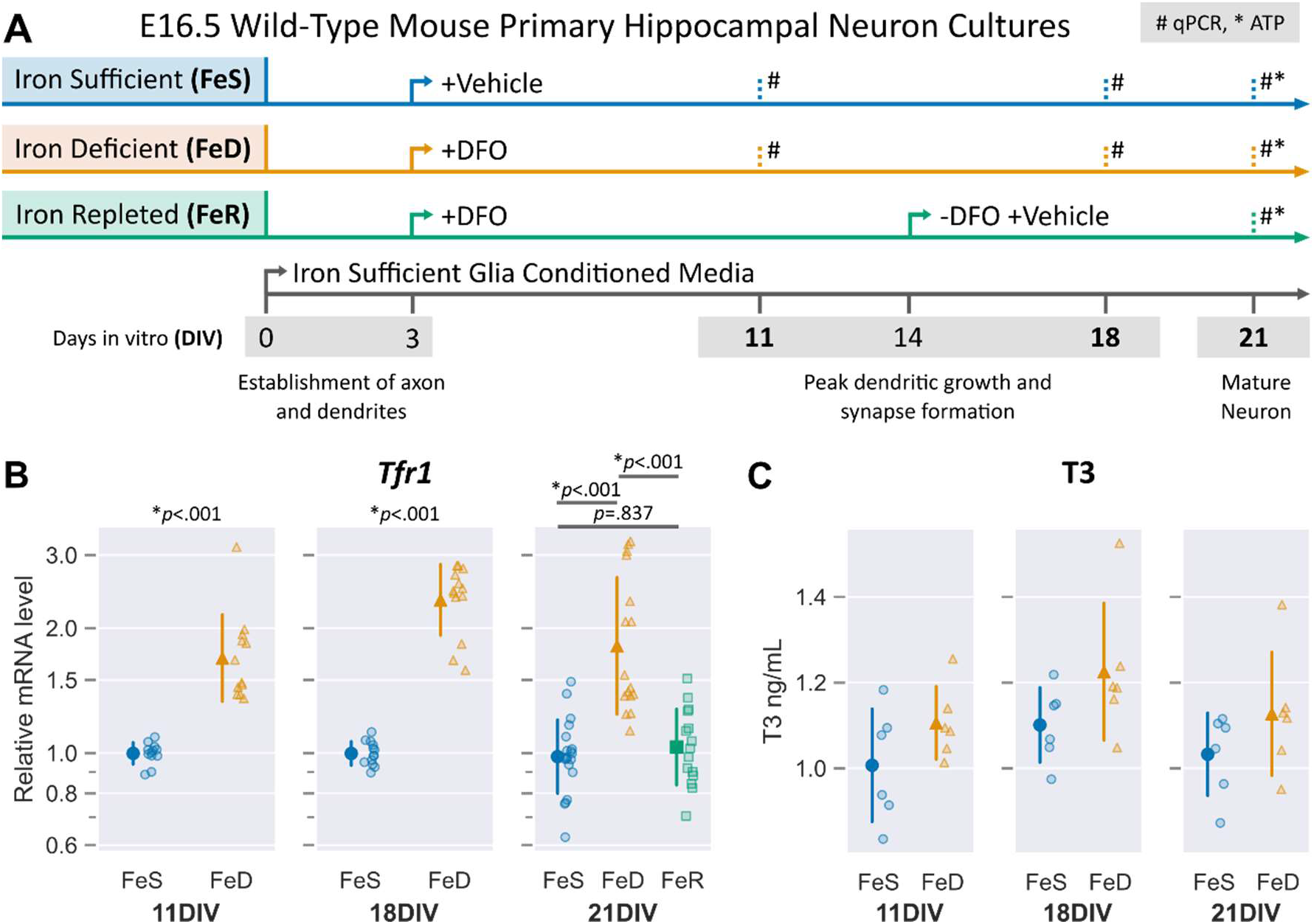
Experimental design of treatment groups and *Tfr1* mRNA expression in developing neuronal cultures. **(A)** The general timeline of key neuron developmental processes from 0 to 21 days of *in vitro* (DIV) culture after harvesting and plating cells from embryonic day 16.5 (E16.5) wild-type mouse hippocampus. Treatment groups are indicated and show initial timing of continual treatment with either vehicle (iron sufficient; FeS) or the iron chelator deferoxamine (DFO) (iron deficient; FeD) in iron sufficient glia conditioned media. Iron repletion (FeR) between 14 and 21DIV was accomplished by removal of DFO. Quantitative real-time PCR (qPCR, #) of selected genes was compared between treatment groups at indicated timepoints; ATP concentration (*) compared between all treatment groups at 21DIV. **(B)** qPCR was performed for *Tfr1,* the gene coding for the main neuronal iron uptake protein, to validate iron status of treatment groups at 11, 18, and 21DIV. Treatment of cultures with DFO increased *Tfr1* mRNA (11, 18, and 21DIV) and could be restored with removal of DFO (21DIV). Individual data points representing one unique culture are shown alongside mean ± SD relative to the average mRNA level to the FeS group at each neuronal age. Relative data are shown on a log scale to accurately reflect the magnitude of changes. Asterisks indicate statistical difference between groups at a given age by Student’s test at 11DIV (*n*=12) and 18DIV (*n*=12-13) and by one-way ANOVA with a Tukey’s post-hoc test at 21DIV (*n*=15-20). **(C)** Extracellular T3 concentrations in culture medium were compared between treatments and across time. A two-way ANOVA found a significant main effect for FeS compared to FeD (F= 6.92, *p* = .013). Extracellular T3 levels were significantly increased in FeD compared to FeS neurons (F = 6.92, *p* = .013), but there was no significant interaction with DIV,

### Iron repletion

For one study, iron repletion was initiated at 14DIV by removing all medium from a subset of DFO-treated cultures and immediately replacing with a half-volume of medium that was removed from iron-sufficient cultures as described (22). A half-volume of fresh glia-conditioned medium containing 5-FU and 0μM DFO was added to bring the medium back to the full volume. The cultures for the iron-sufficient and -deficient groups were subjected to a “mock repletion” as described (22).

### mRNA expression analysis

Cultures were analyzed for thyroid hormone-responsive gene expression just after the beginning of dendritic branching (i.e., 11DIV), during peak branching and synapse formation (i.e., 18DIV), and near maturity (i.e., 21DIV) (Figure 1) (24). Six to 20 independent hippocampal culture vessels per group were used (an independent neurodevelopmental environment is created in each well/vessel (22)) from two to six unique culture preparations. Total RNA was extracted using a Quick-RNA MicroPrep kit (Zymo Research), NucleoSpin RNA XS kit (Macherey-Nagel; Düren, Germany), or TRIzol Reagent with phenol-chloroform extraction (Thermo-Fisher Scientific; Waltham, MA) according to the manufacturers’ protocols as described (20–22). On-column genomic DNA removal was employed for each kit. RNA integrity and purity was established spectrophotometrically using a Nanodrop spectrophotometer. cDNA was synthesized from 100-500 ng total RNA using a High Capacity RNA-to-cDNA Kit (Applied Biosystems). Quantitative real-time polymerase chain reaction (qPCR) was performed using a FastStart Universal Probe Master (Rox) kit (Roche Applied Science) or Luna Universal Probe qPCR Master Mix (New England Biolabs; Ipswich, MA) and a Stratagene MX3000P or Applied Biosystems QuantStudio3 qPCR machine as previously described (20–22). All analyzed samples for each gene had equivalent PCR efficiencies. TaqMan qPCR probes for the genes assessed are outlined in Table 1. Relative mRNA levels for the genes of interest were calculated relative to a reference gene, *TATA box binding protein*, *Tbp*, and are presented as a fraction of the average mRNA level for the iron-sufficient group. We have previously shown *Tbp* to be a stable reference gene for developing iron-sufficient and -deficient hippocampal neuron cultures (20–22).

### Neuronal Culture Media Thyroid Hormone Concentrations

Neuronal culture medium samples were collected at 11, 18, and 21 DIV from iron sufficient and iron deficient cultures (6 media samples/group). T3 and T4 concentrations in the media were measured according to T3 and T4 Total ELISA kit procedures (DRG International; San Francisco, CA). The absorbance of media samples was read at 450nm using the BioTek Synergy LX multimode reader (Agilent; Santa Clara, CA) and concentrations calculated against a standard curve with the Biotek Gen5 software (v3.09).

### Neuronal ATP Concentrations

Intracellular ATP concentrations were measured in 38-45 independent wells per group from two unique culture preparations using the CellTiter-Glo 2.0 reagent (Promega Corp.; Madison, WI) as described (21). ATP concentrations were normalized to the total DNA content of each well as measured using the Quant-iT PicoGreen reagent (Thermo-Fisher; Waltham, MA) as described (21).

### Statistical Analysis

Statistical analyses and data visualization were carried out using Python libraries: numpy (25), pandas (26, 27), seaborn (28), pingouin (29), statsmodels (30), scikit-learn (31), and matplotlib (32). Relative qPCR and ATP data were log-transformed prior to univariate statistical analysis. Student’s t-test (11 and 18DIV qPCR data) or one-way ANOVA with Tukey’s post-hoc multiple comparison test (21DIV qPCR and ATP data), with α = 0.05, was used to determine differences between experimental groups. The untransformed relative data are graphed as mean ± standard deviation (SD) on a log scale to properly visualize the magnitude of changes in both directions. Thyroid hormone concentrations were compared using a two-way ANOVA with DIV and experimental group as between group comparisons; post-hoc tests were not required for this analysis. T3 concentrations (ng/mL) are graphed as mean ± standard deviation (SD).

Principal Component Analysis (PCA) was used to reduce the dimensionality of mRNA relationships. Relative qPCR data was log-transformed and missing values were imputed with the median using a multi-variate between experimental treatment group method using pandas and numpy. PCA was performed using scikit-learn for the maximum number of components and a loadings table and scree plot were generated for all principal components with seaborn. Bi-plots showing the first two Principal Components and loading values for each gene were generated with seaborn; notation of experimental groups on bi-plots were done post-test and did not influence dimensionality reduction.

## Results

Iron sufficient (FeS) primary hippocampal neuronal cultures were compared to iron deficient (FeD) primary neuronal cultures; functional FeD neuronal cultures was accomplished by treating cells at 3DIV with the iron chelator deferoxamine (DFO) (Figure 1A). Cellular iron status was determined by comparing expression of *Tfr1*, which codes for the predominant cell surface protein for neuronal iron uptake, between FeS and FeD cultures (33). *Tfr1* mRNA levels were significantly increased in FeD neurons at 11, 18, and 21DIV (Figure 1B), confirming a functionally iron deficient neuronal state. Extracellular T3 and T4 levels were compared between FeS and FeD cultures across 11, 18, and 21DIV. Extracellular T4 concentration was below the detection limit, as expected since the culture medium does not contain T4. Extracellular T3 levels were significantly increased in FeD compared to FeS neurons (F = 6.92, *p* = .013), but there was no significant interaction with DIV, suggesting that iron deficiency changes T3 handling *in vitro* (Figure 1C).

To investigate if functional iron deficiency in neuronal cultures induces changes in thyroid hormone-regulated gene expression, mRNA levels for genes involved in thyroid hormone homeostasis (*Slc16a2, Slco1c1, Dio2, Crym, Hr*) and neurodevelopment (*Nrgn, Pvalb, Klf9*) were compared between FeS and FeD cultures. At 11DIV, FeD cultures have disrupted thyroid hormone related gene expression as *Hr* (indication of dysregulated thyroid hormone transcriptional control) and *Crym* levels were significantly decreased compared to FeS (Figure 2A). Furthermore, early neuronal differentiation and maturation genes *Klf9, Pvalb,* and *Nrgn* are significantly decreased in FeD cultures compared to FeS cultures (Figure 2B).

**Figure 2.**
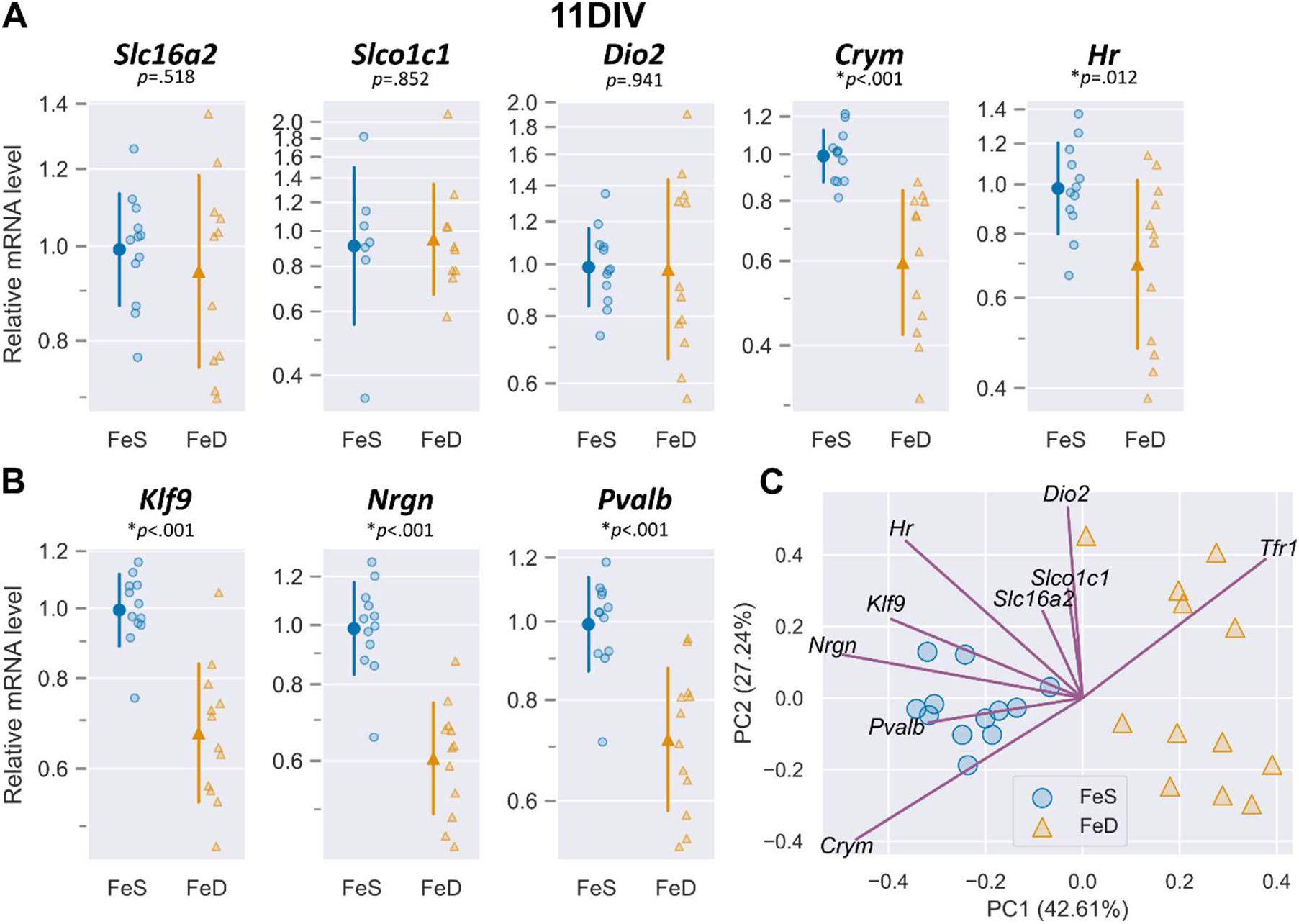
Neuronal iron deficiency alters mRNA expression of thyroid hormone- response genes as early as 11DIV. **(A)** Quantitative real-time PCR (qPCR) was performed for known thyroid hormone homeostasis genes, *Slc16a2, Slco1c1, Dio2, Crym,* and *Hr.* mRNA levels for *Crym* and *Hr* are significantly decreased by iron deficiency (FeD) compared to iron sufficient (FeS) cultures at 11DIV **(B)** qPCR for known thyroid hormone responsive genes involved in neuronal development, *Klf9, Nrgn*, and *Pvalb*. FeD cultures have significantly decreased mRNA levels of these neurodevelopmental genes compared to FeS cultures. For *A* and *B*, individual data points representing one unique culture are shown alongside mean ± SD relative to the average mRNA level for the iron sufficient (FeS) group at each neuronal age. Asterisks indicate statistical difference between groups at a given age by Student’s test (*n*=12). **(C)** Principal component analysis (PCA) plot of mRNA levels for all genes in *A* and *B* and *Tfr1* from Figure 1B. Individual points represent the PCA score for an individual culture and lines represent the loading values for the annotated mRNA. Treatment groups are notated on individual points and reveal clustering of FeS cultures away from FeD neurons on principal component 1 (PC1); FeD neurons have greater spread along PC2. Directionality of loading values suggests that neurodevelopmental genes are more predictive for iron status (*Tfr1*) than thyroid homeostasis genes, with the exception of *Crym*.

At 18DIV, thyroid hormone homeostasis was further disrupted by iron deficiency compared to 11DIV. *Hr* expression was now increased in FeD cultures compared to FeS, suggesting a switch in thyroid hormone transcriptional control (Figure 3A). *Crym* expression remained lower in FeD cultures, but *Slco1c1, Slc16a12,* and *Dio2* expression was significantly increased compared to FeS at 18DIV (Figure 3A). Additionally, FeD cultures normalized expression of the neuronal differentiation gene *Klf9* but maturation genes *Nrgn* and *Pvalb* remained decreased compared to FeS (Figure 3B), indicative of the more mature status of the older, cultured neurons.

**Figure 3.**
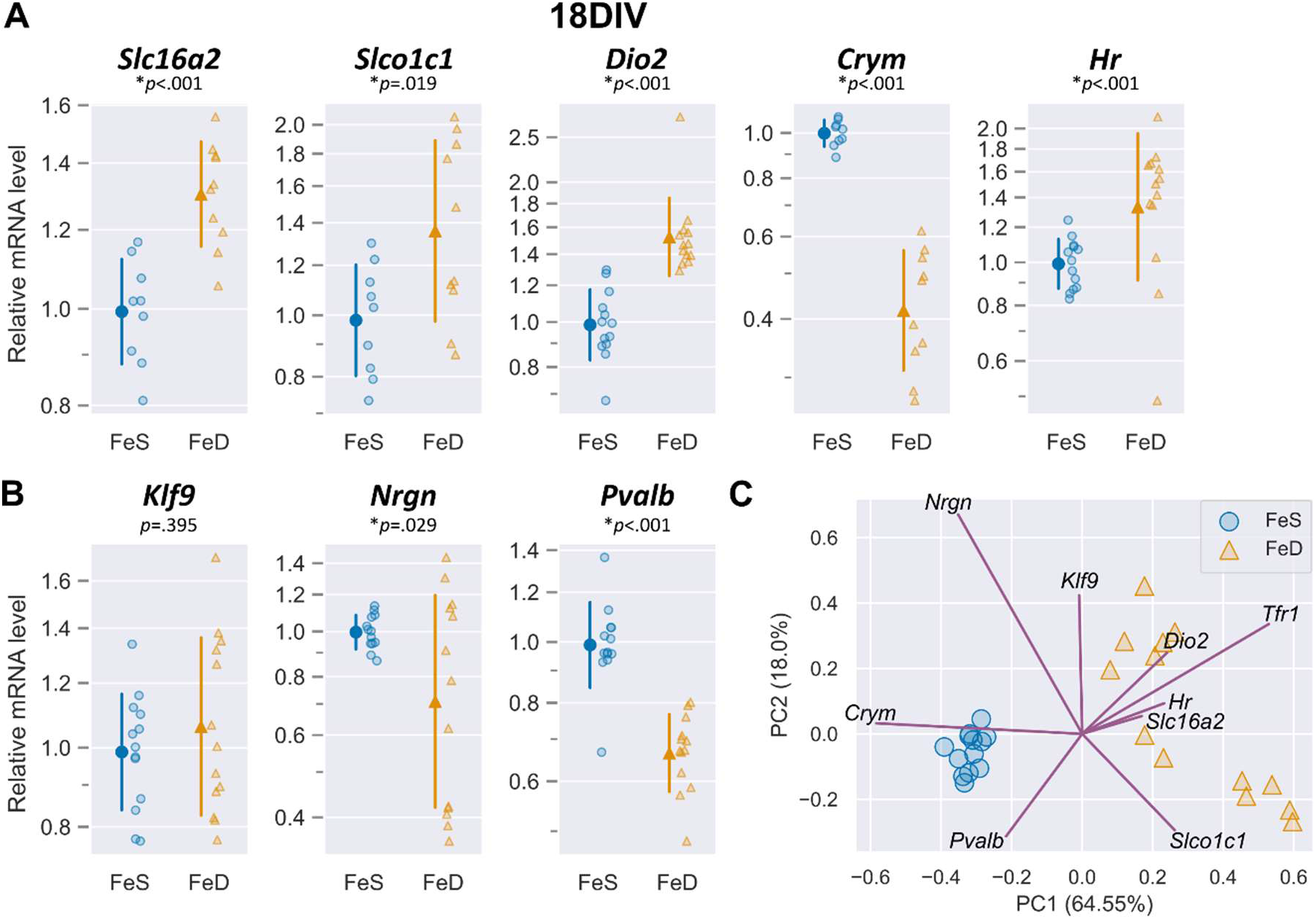
Neuronal iron deficiency alters mRNA levels of thyroid hormone homeostatic genes and thyroid hormone responsive neurodevelopmental genes. **(A)** Quantitative real-time PCR (qPCR) was performed for known thyroid hormone homeostasis genes, *Slc16a2, Slco1c1, Dio2, Crym,* and *Hr.* mRNA levels for *Slc16a2, Slco1c1, Dio2, and Hr* are all significantly increased in iron deficient (FeD) cultures compared to iron sufficient (FeS) cultures, while *Crym* remains significantly decreased as at 11DIV, suggesting cellular sensing of a hypothyroid state. **(B)** qPCR for known thyroid hormone responsive genes involved in neuronal development, *Klf9, Nrgn*, and *Pvalb*. FeD cultures have significantly decreased mRNA levels of *Nrgn* and *Pvalb* compared to FeS cultures. For *A* and *B*, individual data points representing one unique culture are shown alongside mean ± SD relative to the average mRNA level for the iron sufficient (FeS) group at each neuronal age. Asterisks indicate statistical difference between groups at a given age by Student’s test (*n*=12-13). **(C)** Principal component analysis (PCA) plot of mRNA levels for all genes in *A* and *B* and *Tfr1* from Figure 1B. Individual points represent the PCA score for an individual culture and lines represent the loading values for the annotated mRNA. Treatment groups are notated on individual points and reveal clustering of FeS cultures away from FeD neurons on principal component 1 (PC1); FeD neurons have greater spread along PC2. Directionality of loading values suggests that thyroid hormone homeostatic genes are predictive for iron status (*Tfr1*).

While changes to gene expression can be observed between FeS and FeD cultures with univariate data analysis, the variation in severity of functional iron deficiency–as indicated by the up to two-fold difference in *Tfr1* expression level between individual FeD cultures at 18DIV and the 1.5 to 3-fold increase in *Tfr1* expression of FeD cultures compared to FeS cultures (Figure 1B)–cannot be compared to the pattern of thyroid hormone response genes. If the severity of iron deficiency predicts the degree of functional hypothyroidism, then a multivariate analysis would reveal patterns in gene expression whereby increasing levels of *Tfr1* would correlate with further increasing thyroid hormone response genes, for which Principal Component Analysis was used to reduce dimensionality and reveal multivariate patterns in gene expression.

At 18DIV, the largest contributing variable to the first Principal Component (PC1), which predicts 64% of overall gene expression variability, is the iron status indicator *Tfr1* (Figure 3C, S1B). Thyroid hormone homeostatic genes–*Slc16a2, Slco1c1, Dio2, Hr*–positively correlated with *Tfr1* expression except for *Crym,* which strongly and negatively correlated with *Tfr1* expression. Overlaying treatment groups onto the PCA bi-plot (Figure 3C) shows that FeS cultures cluster closely together, but FeD cultures are spread out along both PC1 and PC2. This spread in FeD cultures on the bi-plot indicates that there is underlying variability in the correlated expression profile, of which neuronal maturation genes–*Nrgn, Pvalb,* and *Hr*–correlate more strongly with *Tfr1* expression in PC2 (predicting 18% of the overall variability). Due to the relatively small contribution of PC2 to overall variability, this difference within FeD cultures should be considered sparingly. Overall, multivariate analysis suggests that thyroid hormone homeostatic genes are strong predictors of iron treatment, suggesting cellular sensing of a functionally altered thyroid hormone state of iron-deficient neurons.

In comparison at 11DIV, the first two PCs do not contribute as much predictive strength to the overall gene expression pattern, together only accounting for around 70% of the variability (Figure 2C,S1A). At this stage, *Tfr1* does predict underlying iron status as would be expected, but thyroid hormone homeostatic genes are not as correlated in PC1 (predicting 43%), but are instead correlated in PC2 (predicting 27%). PC1 is primarily affected by thyroid hormone-sensitive neuronal maturation genes–*Klf9, Nrgn, Pvalb* and *Hr*–suggesting that at earlier stages, iron deficiency and any associated thyroid hormone response alters neuronal development gene expression programs.

To determine whether the altered thyroid hormone response in developing iron deficient neurons can be restored to normal, some FeD cultures were repleted with iron by removing the iron chelator DFO between 14 and 2DIV (Figure 1A). These iron repleted cultures (FeR) were compared to FeD and FeS cultures at 21DIV; *Tfr1* mRNA levels were 46% lower in FeR cultures compared to the FeD group and were not significantly different from the FeS group, indicating successful restoration of neuronal iron levels (Figure 1B). However, neuronal ATP levels were only partially recovered after iron repletion and remained 18% lower compared to FeS cultures (Figure 4A), indicating a persistent deficit in neuronal energy metabolism. *Hr, Crym, Dio2*, and *Slc16a2* mRNA levels remained significantly different in FeR cultures compared to FeS cultures (Figure 4B), indicating long-term dysregulation of neuronal thyroid hormone homeostasis after recovery from developmental iron deficiency. In FeR cultures, neuronal maturation genes *Nrgn* and *Pvalb* were not significantly different from FeS neurons, despite continued decrease in mRNA level in FeD neurons compared to control (Figure 4C).

**Figure 4.**
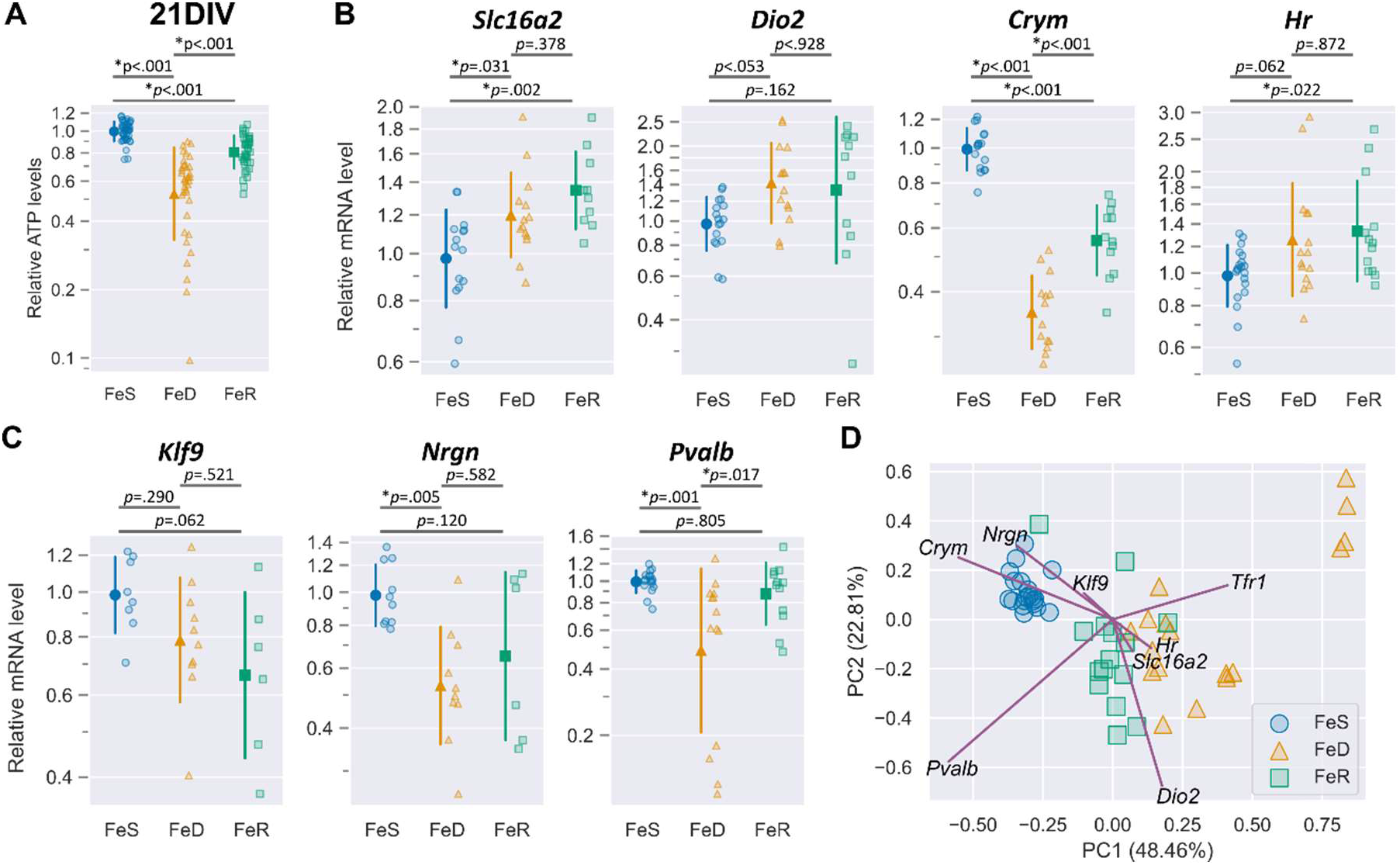
Iron repletion does not fully recover neuronal ATP concentrations nor mRNA levels of thyroid hormone responsive genes at 21DIV. **(A)** Intracellular ATP concentrations were measured and normalized to the DNA content of each well (*n*=38- 45); ATP concentration is reduced in iron deficient (FeD) cultures compared to iron sufficient (FeS) cultures, and is not restored in iron repleted (FeR) cultures (Figure 1A,B). **(B)** Quantitative real-time PCR (qPCR) was performed for known thyroid hormone homeostasis genes, *Slc16a2, Dio2, Crym,* and *Hr.* mRNA levels for *Slc16a2* and *Dio2* are increased in both FeD and FeR cultures relative to FeS cultures. *Crym* is decreased in FeD and FeR cultures, but *Crym* is higher in FeR cultures compared to FeD cultures. *Hr* is not significantly increased in FeD, but is increased in FeR compared to FeS cultures. These changes suggest that repletion of iron status does not restore thyroid hormone homeostatic genes. **(C)** qPCR for known thyroid hormone responsive genes involved in neuronal development, *Klf9, Nrgn*, and *Pvalb*. FeD cultures have significantly decreased mrNA levels of *Nrgn* and *Pvalb* compared to FeS cultures and FeR cultures. Moderate restoration of neurodevelopmental genes with iron repletion suggests some recovery from iron induced changes to thyroid hormone responsive genes. For *A, B,* and *C*, individual data points representing one unique culture are shown alongside mean ± SD relative to the average (A) ATP concentration or (B,C) mRNA level for the iron sufficient (FeS) group at each neuronal age. Asterisks indicate statistical difference between treatment groups at a given age as indicated by Tukey’s post-hoc test after a significant difference between groups via one-way ANOVA (*n*=15-20). **(D)** Principal component analysis (PCA) plot of mRNA levels for all genes in *B, C* and *Tfr1* from Figure 1B. Individual points represent the PCA score for an individual culture and lines represent the loading values for the annotated mRNA. Treatment groups are notated on individual points and reveal clustering of FeS cultures away from FeD cultures on principal component 1 (PC1), with FeR cultures between FeS and FeD cultures, but closer to FeD. Directionality of loading values suggests *Crym*, *Dio2, Nrgn, Klf9, Hr,* and *Slc16a2* are more correlated for predicting variability in cultures than other genes, including the iron status gene *Tfr1*. This suggests that the underlying thyroid hormone responsive gene signature may be useful in predicting current and previous iron status of neuronal cultures.

Again at 21DIV, high mRNA level variability for each gene within treatment groups suggests there may be underlying gene expression correlations (Figure 4B,C). PCA of 21DIV gene expression does not as clearly indicate underlying iron status as indicated by *Tfr1* expression correlating with thyroid hormone response genes (Figure 4D, S1C), because expression of *Tfr1*, but not thyroid hormone-regulated genes, is rescued by iron repletion. Instead, *Crym* and *Pvalb* are the highest predictors of underlying gene variability at 21DIV. Furthermore, the gene expression pattern of thyroid hormone responsive genes–*Slc16a2, Dio2, Hr, Nrgn,* and *Klf9*–correlate most closely with *Crym* at 21DIV. Overlaying treatment groups on the PCA bi-plot (Figure 4D) reveals the multivariate gene expression pattern differentiating FeD and FeR from FeS neurons. The clustering of these treatment groups and predictive pattern of *Crym* and related thyroid hormone responsive genes, but not *Tfr1*, indicates that neuronal cultures repleted with iron maintain a long-term dysregulation of thyroid hormone homeostasis after recovery from developmental iron deficiency.

## Discussion

The major novel finding of this study is that chronic iron deficiency in developing neurons causes impaired thyroid hormone-responsive gene expression indicative of a functionally altered thyroid hormone cellular state, despite normal thyroid hormone availability. This study is the first to demonstrate that neuronal iron status is involved in the regulation of thyroid hormone-target genes in a cell-intrinsic manner. These findings demonstrate a need for future studies of the undiscovered intracellular mechanism coordinating iron and thyroid hormone activities during neuron development since iron-dependent *de novo* thyroid hormone synthesis only occurs in the thyroid gland. An argument can be made for iron and thyroid hormone activities to be closely matched in the developing brain as part of a metabolic adaptation program. Thyroid hormones regulate cellular energy demand in the developing brain through their functional role in stimulating ATP-consuming processes, including generation of ion gradients, cytoskeleton polymerization, and cell proliferation (3). T3 is also a key regulator of nuclear and mitochondrial transcription for cellular energy metabolism genes (e.g., cytochrome c oxidase subunits) and mitochondrial biogenesis (e.g., PGC1-alpha) during brain development (34). Similarly, iron is essential for mitochondrial energy production in the developing brain through its direct role in the structure and redox activity of cytochrome- and iron-sulfur cluster-containing TCA cycle and electron transport chain proteins (4). Brain development involves multiple energetically demanding processes— such as cell proliferation and differentiation, neuronal dendrite/axon outgrowth, synapse formation/function, and myelination—that require both optimal iron and thyroid hormone status (35, 36). Thus, we offer the novel hypothesis that matching the neuronal metabolic and growth rate (mediated by thyroid hormone) with the availability of metabolic substrates for oxidative phosphorylation (e.g., iron) is a part of a key neuroplasticity mechanism that adapts the developing neuron/brain to metabolic disruptions and prevents long-term damage.

In support of this hypothesis, both iron and thyroid hormone regulate the differentiation of Parvalbumin-positive interneurons and the formation of perineuronal nets (37–40), which are key indicators of the opening and closing of critical neurodevelopmental windows, respectively (41). Across the three DIVs we tested, *Pvalb* expression was the most consistently, strongly and negatively correlated gene with iron status (*Tfr1*). Thus, the reduced *Pvalb* expression observed in FeD neurons is potentially part of an adaptive response to delay a critical developmental window because of insufficient metabolic substrate to meet the high energy requirements of this period.

Alternatively, our iron repletion study demonstrates that there is a cost to prolonged developmental iron deficiency, in that there are residual perturbations to thyroid hormone-responsive gene expression and ATP production even after neuronal iron status is restored. Dietary fetal-neonatal iron deficiency in rodents causes reduced circulating and brain thyroid hormone levels and dysregulation of thyroid hormone-responsive gene expression in the neonatal brain (14–16). This *in vivo* disruption to thyroid hormone action in the iron deficient brain is likely driven through a systemic effect, caused by impaired activity of the iron-dependent thyroid peroxidase (TPO) enzyme (11), which is responsible for iodination and tyrosine coupling steps of thyroid hormone synthesis in the thyroid gland. Hu *et al*., showed that dietary maternal iron deficiency impairs neonatal offspring brain thyroid hormone levels even without the brain becoming iron deficient, supporting the hypothesis that this is a downstream result of impaired thyroidal thyroid hormone synthesis (17). Given the critical importance of maintaining normal levels of both iron and thyroid hormone for proper brain development, concurrent alterations in both iron and thyroid hormone status would generate a “double hit” to the developing brain and result in poorer outcomes than either condition alone. Although neuronal iron and thyroid hormone coordination could be adaptive in the short-term, it is likely maladaptive in the long-term if thyroid hormone action is unable to be restored to normal with iron repletion.

Cellular response to thyroid hormone deficiency involves compensatory increases in expression of thyroid hormone transporters and “activating” deiodinases to maintain thyroid hormone homeostasis. We show that 11DIV iron deficiency reduces the expression of classic, positively thyroid hormone-regulated genes (i.e., *Klf9*, *Nrgn*, *Pvalb*, and *Hr*) without an increase in expression of negatively regulated thyroid hormone transporter (i.e., *Slc16a2* and *Slco1c1*) and deiodinase (i.e., *Dio2*) genes. This suggests that it is not a lack of intracellular T3 availability that is causing decreased expression of these thyroid hormone-regulated genes since that would also result in compensatory increases in expression of thyroid hormone import and activation genes. At 18DIV, however, *Slc16a2, Slco1c1*, and *Dio2* mRNA levels are increased, suggesting that the iron deficient neurons are sensing low T3 availability at this stage. Since neurons do not synthesize thyroid hormones, iron deficient neuron cultures have normal thyroid hormone availability in the medium. These findings raise questions of what happens to the “unused” T3 in iron deficient cultures and why these neurons are seemingly sensing low T3 availability. Our data show a slight, but significant, increase in extracellular T3 concentration in iron-deficient neuron cultures suggesting decreased T3 import or increased export. SLC16A2 *(Slc16a2*/Mct8) has been shown to also function as a T3 exporter (42); thus, increased expression of *Slc16a2* as a result of iron deficiency may instead be indicative of a cellular response to decrease the intracellular availability of T3, in order to match the decreased metabolic potential of available iron and prevent oxidative stress.

As further evidence for iron deficiency driving T3 export, the second highest predictor of underlying gene variability between iron sufficient and iron deficient neurons at both 11 and 18DIV was *Crym*, a T3 binding protein. *Crym* expression is negatively correlated in both PC1 and PC2 with *Tfr1* expression at these stages. At 21DIV, gene variability was more strongly predicted by *Crym* than *Tfr1*, suggesting that cultures replete with iron maintain a signature of gene expression (i.e. *Crym*) indicative of previous iron status. Decreased *Crym* expression has been shown to increase T3 efflux through SLC16A2 and simultaneously decrease nuclear thyroid hormone activity (43). Since neuronal iron deficiency decreases *Crym* expression and increases *Slc16a2* expression, this provides further support that low neuronal iron status may result in the cell increasing T3 efflux to reduce overall thyroid hormone function, including genomic and non genomic functions, including mitochondrial metabolism and cytos keletal dynamics required to grow developing neurons (44). To understand the cellular handling of dysregulated thyroid hormone function and availability, future studies will need to assess intra- and extra-cellular concentrations of thyroid hormone metabolites, thyroid hormone transport kinetics, and deiodinase activities in iron deficient neurons.

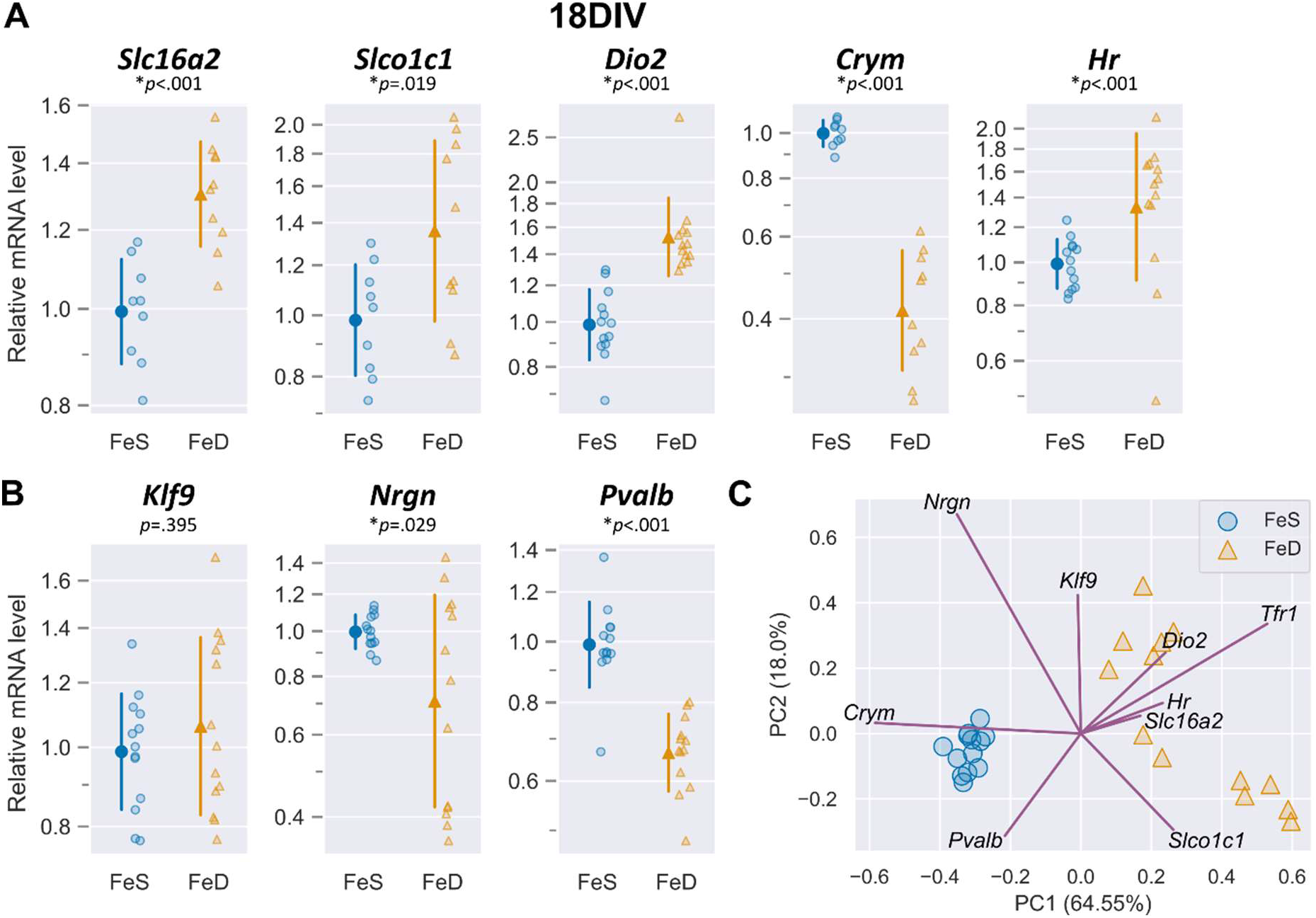

In the neuron, there is no known iron dependent mechanism directly controlling intracellular thyroid hormone levels, availability, or action. However, Hairless (*Hr*) is an interesting candidate protein; *Hr* expression in iron deficient neurons was decreased at 11DIV, increased at 18DIV, and persistently increased at 21DIV even after iron repletion. Since *Hr* gene transcription is directly and positively regulated by T3/Thyroid hormone receptor, this temporal variation without direct changes to thyroid hormone availability suggests additional non-thyroid hormone regulatory control of *Hr* expression in iron deficient neurons. Hairless is also directly involved in thyroid hormone-regulation of transcription through its role as a thyroid hormone receptor corepressor via its histone demethylase activity, which requires an iron-containing Jumonji C (JmjC) domain (33). Thus, thyroid hormone receptor control of thyroid hormone action may be directly regulated by neuronal iron availability through Hairless, a hypothesis which requires further investigation.

If there is an underlying adaptive mechanism mediating the effect of iron deficiency on thyroid hormone action, then there should be a reciprocal effect with thyroid hormone deficiency causing a decrease in iron availability or iron-dependent activity. In our previous studies, mild, moderate, or severe fetal-neonatal thyroid hormone deficiency did not alter the total iron content of the neonatal brain (14–16), but these studies did not measure cellular iron storage, availability, or utilization. In a fetal-neonatal rat model, systemic hypothyroidism increases ferritin L (iron storage) and decreases ferritin H (iron utilization) expression in the developing brain (33). Hyperthyroidism has the opposite effect (33), indicating a direct relationship between brain thyroid hormone availability and the balance of cellular iron storage and utilization. Further investigation into thyroid hormone control of intracellular iron storage, availability, and utilization will be important to understand the regulatory mechanisms of iron-thyroid hormone interactions and their roles in maintaining the balance between metabolic supply and demand during brain development.

Although our findings are from mouse cell cultures, there are potential translational implications. Iron status during pregnancy has been shown to be an independent risk factor for thyroid disorders including subclinical hypothyroidism, thyroid autoimmunity, and maternal hypothyroxinemia (12, 13). Maternal iron deficiency and mild thyroid dysfunction are associated with similar adverse offspring neurological outcomes (e.g., increased risk of cognitive deficits, ADHD, autism, and schizophrenia) (5, 10, 35). Thus, interactions between iron and thyroid functions have consequences for human fetal-neonatal and child brain health, indicating the need to better understand iron and thyroid hormone interactions for proper clinical management of these common early life disorders.

## Acknowledgements

We thank the members of the Bastian, Georgieff, and Lanier labs for their invaluable assistance with culture preparation and data interpretation. Grants supporting this research included NIH R01 HD029421 (MKG), R01 HD094809 (MKG), F32 HD085576 (TWB), T32 HL007062 (TRM) and an American Thyroid Association Research Grant (TWB).

## Author Contributions

TWB, LML, and MKG designed the research; TWB conducted the neuronal culture experiments; TWB, SEB, and KNG conducted the gene expression experiments; EHT conducted and analyzed the thyroid hormone ELISA experiments; TRM, TWB, EHT, SEB, and KNG performed the statistical analyses. TRM and TWB wrote the manuscript and had responsibility for final content. MKG and LML edited the manuscript. All authors read and approved the final manuscript.

## Data Sharing

Data described in the manuscript, code book, and analytic code is publicly and freely available without restriction at https://github.com/TimMonko/2023-FeD-TH-manuscript.

