## Supplemental Figure for "Cellular Iron Deficiency Disrupts Thyroid Hormone Regulated Gene Expression in Developing Hippocampal Neurons"

**Supplement Figure**

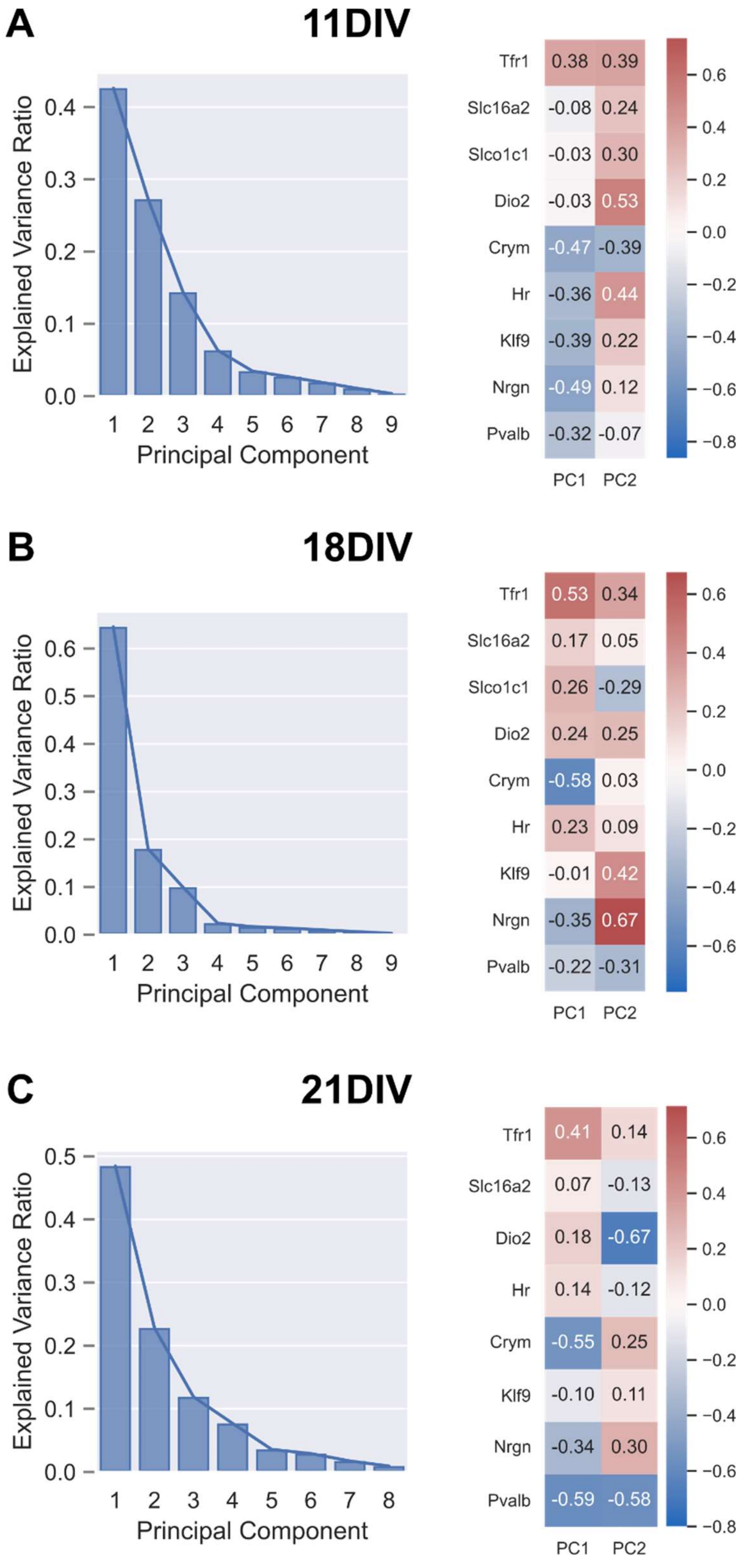

**Supplement Figure 1. Scree plots and loading heatmaps for Principal Component Analyses.** Scree plots showing the proportion of overall variance accounted for by each Principal Component corresponding to the appropriate PCA at each DIV are shown on the left (A: 11DIV / Figure 2C; B: 18DIV / Figure 3C; C: 21DIV / Figure 4D). Corresponding heatmaps of loading values are shown on the right to show the representative weight of each gene's mRNA level prediction to each Principal Component.
